# Does cellular adaptation to force loading rate determine the biphasic vs monotonic response of actin retrograde flow with substrate rigidity?

**DOI:** 10.1101/2021.04.23.441062

**Authors:** Partho Sakha De, Rumi De

## Abstract

The transmission of cytoskeletal forces to the extracellular matrix through focal adhesion complexes is essential for a multitude of biological processes such as cell migration, differentiation, tissue development, cancer progression, among others. During migration, focal adhesions arrest the actin retrograde flow towards the cell interior, allowing the cell front to move forward. Here, we address a puzzling observation of the existence of two distinct phenomena: a biphasic relationship of the retrograde flow and cell traction force with increasing substrate rigidity, with maximum traction force and minimum retrograde flow velocity being present at an optimal substrate stiffness; in contrast, a monotonic relationship between them where the retrograde flow decreases and traction force increases with substrate stiffness. We propose a theoretical model for cell-matrix adhesions at the leading edge of a migrating cell, incorporating a novel approach in force loading rate sensitive binding and reinforcement of focal adhesions assembly and the subsequent force-induced slowing down of actin flow. Our model unravels both biphasic and monotonic responses of the retrograde flow and cell traction force with increasing substrate rigidity, owing to the cell’s ability to sense and adapt to the fast-growing forces. Moreover, we also elucidate how the viscoelastic properties of the substrate regulate these nonlinear responses and alter cellular behaviours.

## Introduction

Living cells interact with the extracellular matrix and other cells through the generation and transmission of forces from the actin cell cytoskeleton to the cell environment [1, 2]. The transmission of forces is crucial for diverse cellular processes such as cell spreading, migration, differentiation, morphogenesis, wound healing, and tissue regeneration [1–5]. Cells form physical linkages known as focal adhesions at the cell-matrix interface and transmit forces to the extracellular matrix. During cell movement, myosin motors pull the actin-filaments towards the cell center in a process known as the retrograde flow. The focal adhesions (FAs) arrest the retrograde flow and slow it down by transmitting the forces to the substrate. This slowing down allows for actin polymerization and cell protrusion to advance at the cell front. The dynamic variation of retrograde flow, orchestrated with the assembly and disassembly of focal adhesions, gives rise to a repetitive jerky motion, known as ‘stick-slip’. Stick-slip motion has also been observed in many passive systems, such as peeling of adhesive tape [6–8], earthquakes [9], pulling wet finger across a glass pane [10], among others. Systems exhibiting stick-slip behaviour, in general, go through the state where the system spends most time in building up energy, known as the ‘stuck’ phase, and then followed by a quick release of the gained energy through the ‘slip’ phase.

Several experiments have demonstrated that substrate rigidity plays a crucial role in driving cellular adhesion, spreading, and migration processes. However, a debate exists on how the cell traction force or actin flow is regulated with the variations in the substrate stiffness. On the one hand, observations from a group of experiments on embryonic chick forebrain neurons and glioma cells, have revealed neither very soft nor very stiff substrates are beneficial for efficient cell migration; the optimal stiffness lies somewhere in between, where the focal adhesions able to most efficiently transmit cytoskeletal forces to the substrate [11, 12]. In these experiments, it is observed that traction force and retrograde flow velocity exhibit a biphasic behaviour with increasing substrate rigidity with the presence of an optimal substrate stiffness corresponding to minimal retrograde flow and maximal traction force. Moreover, experiments performed on neutrophils [13] and smooth muscle cells [14] also show this biphasic relationship. On the contrary, many experimental studies show a monotonic relationship of traction force with the variation in substrate stiffness, where traction force increases with the increase in substrate stiffness [15–18]. Interestingly, recent experiments performed on mouse embryonic fibroblasts suggest that the dynamic mechanism of the molecular clutch that connects actin to the extracellular matrix and transmits forces is regulated by the force loading rate, which varies with substrate rigidity. Rate-dependent unfolding and binding of the adapter proteins of the molecular clutch, particularly the protein talin, can account for the biphasic and monotonic behaviours of traction force with increasing substrate rigidity [2, 19].

There have been many attempts at developing theoretical models to understand the cell migration process and the strong dependence on substrate stiffness. For example, a mechanical model was developed by Barnhart *et al*., to address periodic oscillations of the cell length caused by alternating stick-slip cycles at the cell edge of motile fish epithelial keratocytes [20]. Using an analytical formulation of the molecular clutch model, Sens has shown ‘stick-slip’ dynamics of cell-traction could explain periodic waves of protrusion/retraction and propagating waves along the cell edge, leading to steady crawling or bipedal motion, and bistability [21]. Weinberg *et al*., developed a mechanical model which described traction force generation from the cell and subsequent assembly of the matrix protein fibronectin, which is crucial for cellular mechanical response [22]. Moreover, stochastic models for stick-slip and smooth migration modes for polarised cells have also been proposed [23]. Chan and Odde have modelled cell-substrate adhesions as stochastically binding and unbinding molecular clutches connecting the actin cytoskeleton to the substrate [11]. Through this model, they were able to exhibit the experimentally observed stick-slip dynamics for filopodial growth cones as well as the biphasic relationship of traction force and retrograde flow velocity with substrate stiffness. Further studies based on the model provided more insight into how different parameters influence the dynamics and the presence of different optimal stiffness for different cell lines [12, 24]. Other models involving stochastic bond dynamics, which integrated traction force-dependent retrograde actin flow, could also capture the biphasic force-velocity relation [25–27]. On the other hand, in the studies conducted by Elosegui-Artola *et al*., the presence of two different type integrin molecules, namely *α*_5_*²*_1_ and *α_υ_β*_6_, expressed for healthy and malignant tissues, with different binding and dissociation rates were considered. This, combined with increased ligand affinity when the bond force increases beyond a certain threshold, could explain their observation of increasing traction force and decreased retrograde flow velocity at higher substrate stiffness [28]. In another model, considering talin unfolding above a threshold rigidity also produced similar results [2, 19]. Furthermore, Gong *et al*. [29] have studied how the substrate rigidity regulates the biphasic and monotonic behaviours of cell spreading involving a FA reinforcement mechanism where the binding rate changes above a cut off force. We have also investigated how the viscoelastic properties of the substrate affect the stick-slip dynamics and the biphasic response of migrating cells [30]. Even though there has been a significant amount of theoretical work done on this topic, there still exists a lack of a single encompassing model where both of these starkly different cellular responses - biphasic versus monotonic behaviour of actin retrograde flow and cell traction force dynamically emerge with the variation in viscoelasticity of the substrate.

In this work, we propose a theoretical model for the stick-slip dynamics at the cell leading edge to investigate the nonlinear response of biphasic vs monotonic behaviour with varying substrate rigidity. It follows the formulation of coupled reaction-diffusion equations involving the known key players in cell protrusion and motility, such as actin flow, myosin motors, focal adhesions integrated with viscoelasticity of the cell-matrix system. The crux of the model lies in the force loading rate sensitive binding and reinforcement of focal adhesions assembly that could predict the experimentally observed puzzle of biphasic vs monotonic response of actin retrograde flow and cell traction force with increasing substrate stiffness. The difference in the cellular behaviours is attributed to cells ability to sense the force loading rate and adapt to it through the recruitment of new adhesion molecules to balance the fast-growing forces. Notably, both biphasic and monotonic responses of cell traction force and actin flow dynamically emerge from our model without the need for imposing a cut-off force or a threshold value as considered by earlier studies. Our theory further elucidates how the substrate viscosity along with substrate elasticity alter these nonlinear cellular responses. Besides, it also predicts the loss of cell sensitivity to differentiate between soft and stiff substrates when the substrate viscosity is high and vice versa.

## Model and Methods

The model has been developed based on our previous work [30] of transmission of force from the cell actin cytoskeleton to the extracellular matrix (ECM) through the formation of physical linkages with the connector proteins providing a dynamic link between the cell and ECM. The model consists of free receptors diffusing in the cell cytoplasm, which bind with the ligands on the substrate to form bound receptors or cell-substrate adhesion bonds as illustrated in Figure 1. These adhesions are modelled as molecular clutches which transmit forces from the cell cytoskeleton to the ECM. They also serve to slow down the retrograde flow of actin, thus allowing further protrusion at the cell front. Under the effect of growing force, as the F-actin coupled to the molecular clutches is pulled by the myosin motors, these adhesions disintegrate and revert back to free receptors and ligands. The spatio-temporal evolution of the densities of the free and bound receptors are given by the following coupled reaction-diffusion equations,

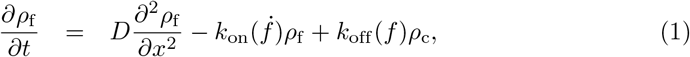

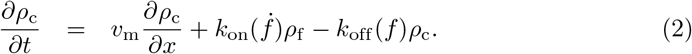

**Figure 1.**
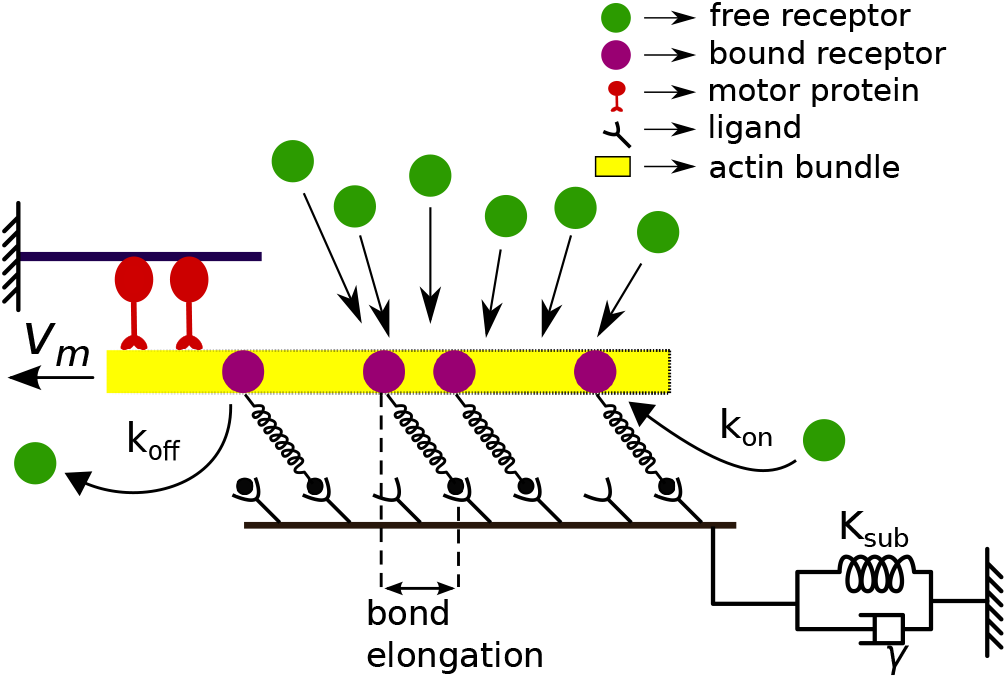
Schematic of the model: Free receptors (green circles) diffusing in the cytoplasm bind with the ligands on the substrate to form bound receptors or closed bonds (magenta circles), which act as the physical linkage between the cell and the substrate. The myosin motors pull on the F-actin, flowing inwards with the retrograde velocity, *υ_m_*, slowed by the adhesion bonds. The viscoelastic substrate is represented by a Kelvin-Voigt system of a spring and a viscous damper.

Here *ρ*_f_ and *ρ*_c_ denote the densities of free receptors and bound receptors respectively, *D* is the diffusion constant for free receptors and *υ_m_* is the actin retrograde flow velocity. The total number of free and bound receptors is taken to be conserved, 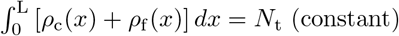. The ligand-receptor adhesion bonds are modelled as Hookean springs with stiffness *K*_c_. Thus the force, *f*, on an individual bond is calculated by multiplying *K*_c_ with the elongation of the spring. Further, *k*_on_ and *k*_off_ describe the binding and unbinding rates of free and bound receptors, respectively.

It has been observed in several experiments that mechanical forces significantly affect the growth and stability of focal adhesions [31, 32]. For example, the presence of time-varying stretches, reorganize and reorient FAs, actin cytoskeleton, and thus alter the response of the cell [33, 34]. Recent studies show that the force loading rate is another crucial determining factor for the mechanosensitivity of adhesion clutches that connect actin to the ECM and regulate the force transmission [2]. It is observed that the ligand-receptor binding affinity, unfolding the adapter proteins in the molecular clutch, and subsequent focal adhesions growth is modulated by the force loading rate [19]. Further, studies on bond force spectroscopy indicate that bond lifetime and rupture-force exhibit an adhesion reinforcement with increasing loading rate [35, 36]. Following the experimental findings, we consider the binding rate of ligand-receptor adhesion bond formation to be dependent on the force loading rate. In our model, the binding rate is proposed to have a reinforcement mechanism as, 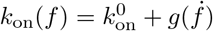, where 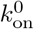 is the unloaded binding rate and 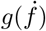 is the loading rate dependent binding. For simplicity, we assume a linear form, 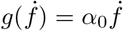, where the binding factor, *α*_0_, denotes the tendency of the cell to sense the growing force and adapt to it through the formation of new bonds to strengthen the focal adhesions cluster. Moreover, we study the dynamics of the focal adhesions cluster exhibiting both catch bonds and slip bonds behaviours and the dissociation rate is expressed in the form of 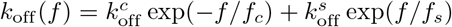 for catch bonds and 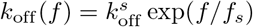 for slip bonds; where 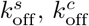 are unloaded slip dissociation rate and unloaded catch dissociation rate respectively [37]. While, *f_s_, f_c_* are the characteristic slip and catch bond dissociation forces.

Now, the equation of motion for the viscoelastic substrate is obtained by balancing the viscous and the elastic forces arising from the substrate with the total force experienced by all adhesion bonds as,

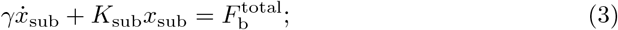

where, *γ* represents the substrate viscosity, *K*_sub_ is the substrate elastic stiffness, and 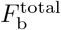 is the total traction force, given as, 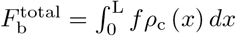. *L* is the spatial extent of the adhesion patch. The slowing down of retrograde flow due to the traction force is given by, 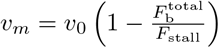, where *υ*_0_ is the unloaded velocity. Here, *F*_stall_ is the stall force due to the pull of the myosin motors and is expressed as *F*_stall_ = *n_m_* * *F_m_*, where *n_m_* is the total number of myosin motors present and *F_m_* is the force exerted by a myosin motor.

Migrating to a dimensionless formulation, we represent the scaled space coordinate as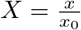, where 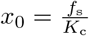. The dimensionless time is given by the form *τ* = *k*_0_*t*. The scaled densities of the free and bound receptors are given as 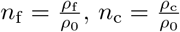, where *ρ*_0_ denotes the average density of the receptors and is defined as, 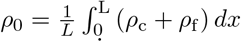. The dimensionless association and dissociation rates are, 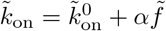 and 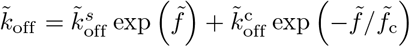, where 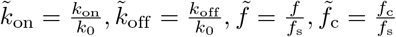, and *α* = *α*_0_*f*_s_. Other dimensionless variables are expressed as: 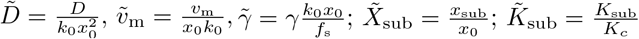, and 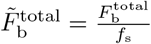. Thus the scaled equations of motion are given as,

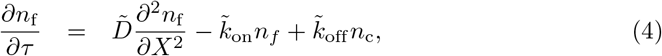

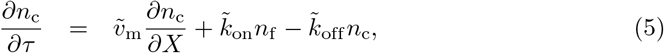

and

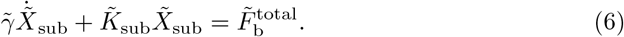

## Results

We investigate the dynamics by solving the reaction-diffusion equations coupled with the equation of motion of the substrate, Eqs. 4-6 numerically. The spatial component is solved through discretization by finite difference method on a grid size *N* and the time evolution is studied using a fourth order Runge-Kutta method. The boundary conditions are considered to be such that the total number of receptors: both free and bound receptors are conserved. The bound receptors do not diffuse and the influx of free receptors at the adhesion patch is assumed to be same as the outflux. Here we present results for catch bonds considering a representative set of parameters inspired from real experimental values, such as 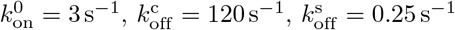, *f*_c_ = 0.5 pN and *f*_s_ = 1pN [37]. The unloaded actin retrograde flow velocity is taken as *υ*_0_ = 120nms^−1^, myosin stall force, *F*_m_ = 2pN, and number of myosin motors, *nm* = 100. The diffusion constant is considered as 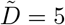. The range of substrate elastic stiffness and substrate viscosity are also inspired from experimental values as *K*_sub_ ~ 0.01 – 1000 pN nm^−1^ [11, 24], and the range of substrate viscosity is *γ* ~ 0.01 – 100pN.snm^−1^ [38].

Figures 2(a) - (d) show the time evolution of the total traction force, 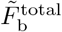, on soft and stiff substrates for different loading rate-sensitive binding affinity, *α*, of ligand-receptor adhesion bonds. In the case of soft substrates shown in Figs. 2(a)-(b), since the substrate is very compliant, the build-up of bond force, 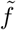, is also very slow. Thus, the traction force slowly increases over time and in turn reduces the retrograde flow of actin, 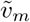 (shown in supplementary) allowing actin polymerization to advance at the cell leading edge, giving rise to the ‘stuck’ state. With an increase in the bond force, the dissociation rate first decreases due to the catch bond behaviour and then starts increasing after a certain force value. Thus, the binding rate can no longer keep up with the exponentially growing dissociation rate, and the adhesion cluster starts to dissociate rapidly. As a result, the traction force also decreases fast; as there is nothing to anchor to the substrate, the actin filament slips backwards, and the retrograde velocity increases rapidly, giving rise to the ‘slip’ state, and the stick-slip cycle continues. Comparing Figures 2(a) and 2(b) on soft substrates, we see there is no significant difference in the nature of the evolution of the traction force with varying *α*. Since soft substrates deform easily, the force on individual bonds slowly builds up with time, resulting in a low force loading rate. Therefore, the binding reinforcement factor, *α*, doesn’t make much of a noticeable difference in the dynamics. However, the situation changes remarkably when we migrate to a stiff substrate, as seen from Figures 2(c) and 2(d). With the increase in substrate stiffness, we see an increase in the force loading rate due to the rapid build-up of force on the adhesion bonds since the stiffer substrate does not deform easily. Here, we find a marked difference in the evolution of traction force due to different loading rate-dependent binding rates for different cell types. Cells having a stronger force loading rate sensitive binding affinity, *i.e*., with a higher value of *α*, as the binding rate increases with the growing loading rate, more free receptors bind with the substrate ligands, and thus, the adhesion gets strengthen. As a result, the total traction force increases (Fig. 2 d), which in turn reduces the retrograde flow velocity, and the stick-slip duration also increases. On the contrary, with a low value of *α*, cells that are not very sensitive to the force loading rate exhibit lower traction force and shorter stick-slip cycles on the stiff substrate as shown in Fig. 2 c. Dissociation rate dominates due to the fast-rising force that shortened the lifetime of the bonds, and the cell is unable to transmit the force to the substrate efficiently.

**Figure 2.**
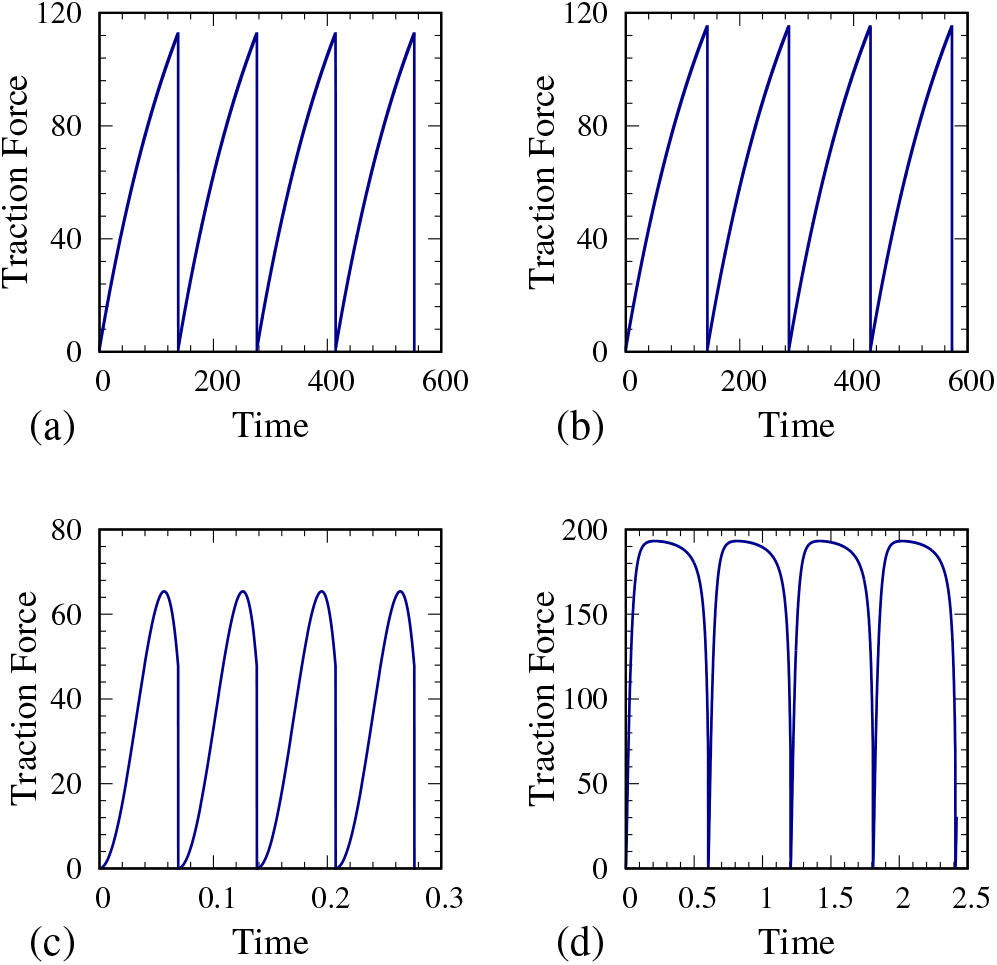
Time evolution of the traction force, 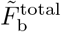, on soft and stiff substrates for two different values of the loading rate dependent binding affinity, *α*; time evolution on soft substrate shown in (a) 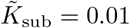, *α* = 0.01 and (b)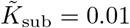, *α* = 1; whereas on stiff substrate in (c) 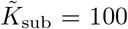, *α* = 0.01 and (d) 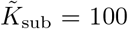, *α* =1. (Keeping 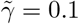 to a constant value.)

Figures 3*a* and 3*b* show the average value of the traction force, 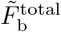, and the retrograde velocity, 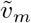, with increase in substrate elastic stiffness, 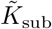, for different values of cellular force loading rate sensitive binding affinity, *α*. As the binding factor *α* increases, a clear transition is seen in the nature of variation of traction force and retrograde flow at higher substrate stiffness. For low α values, we observe a biphasic relationship and the existence of an optimum substrate stiffness with maximum traction force and minimum retrograde flow. In such cases, cells do not have any binding reinforcement mechanism with higher loading rates on stiffer substrates; therefore, the fast-growing dissociation rate dominates over the binding rate, and the bonds break off quickly, resulting in low traction force and high retrograde flow velocity. Now, as the substrate stiffness is lowered, substrate starts deforming easily, and the bond force becomes low enough for the cell to form substantial adhesion linkages, which enable the efficient transmission of force from the actin cytoskeleton to the substrate, thus increasing the traction force and lowering the retrograde flow. However, if the substrate is made even softer since the individual bond force grows very slowly, they spend a large amount of time experiencing low forces; this lowers the average value of the traction force and increases the retrograde flow. All of these combined give rise to the biphasic relationship with increasing substrate rigidity. On the other hand, with higher values of *α*, *i.e.*, when the cell can sense high force loading rate and adapt through increased ligand-receptor binding affinity to balance the fast-growing forces, the biphasic relationship disappears, and a monotonic increase of traction force and monotonic decrease of retrograde flow velocity is observed. The effect of cell types having a stronger dependence on loading rate becomes prominent on the stiffer substrate where the force loading rate is very high. Here, the binding rate significantly increases due to the loading rate dependent reinforcement; more free receptors are recruited to form new bonds, strengthen the adhesion, and allow more efficient transmission of cytoskeletal force to the stiffer substrate. This leads to a monotonic increase of traction force and consecutively monotonic decrease of retrograde flow as the substrate gets stiffer as shown in Figures 3*a* and 3*b*.

**Figure 3.**
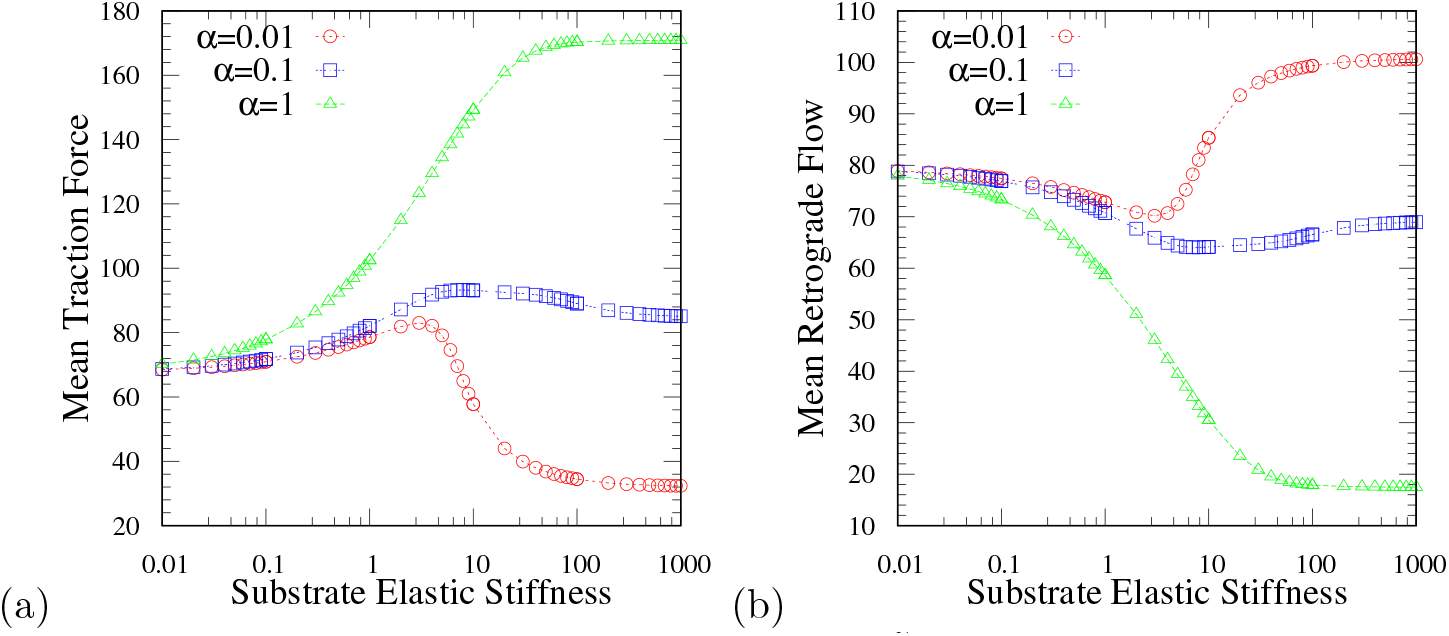
Average value of (a) the traction force, 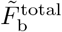, and (b) the retrograde flow velocity, 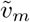 exhibiting the transition from the biphasic to the monotonic behaviour with increasing substrate elastic stiffness, 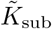, and with varying values of the loading rate dependent reinforcement factor, *α* = 0.01 (red circle), 0.1 (blue square), 1 (green triangle); keeping 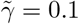.

Substrate viscosity also plays an important role in determining the cellular response, how cells spread, migrate, and transmit forces to the extracellular matrix [29, 38]. Here, we investigate how the substrate viscosity affects the biphasic and monotonic behaviours of actin retrograde flow and the traction force. In Figure 4, from the contour plots of the average traction force and average retrograde velocity, we can distinctly differentiate how substrate viscosity along with substrate elasticity influences cellular response for two cases: when cells do not have any mechanism to adapt to the increasing force loading and where cells strengthen the adhesions by forming new ligand-receptor bonds to balance a rapidly growing force represented by a low and high value of *α* respectively. As shown in Figures 4a and 4b, in case of smaller *α*, when the substrate elastic stiffness, 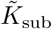, is low, the biphasic behaviour emerges with varying substrate viscosity, 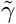. There exists an optimal substrate viscosity for which one observes a maximal traction force and a minimal retrograde flow velocity. Similarly, when the substrate viscosity is low, one observes an optimal substrate elastic stiffness. However, when the substrate viscosity is increased beyond a particular value, not only the biphasic relationship disappears, but the cell also loses its ability to sense variations in substrate elastic stiffness. Likewise, when the substrate elastic stiffness is increased beyond a particular value, the presence of optimal viscosity disappears, and the cell becomes insensitive to variations in substrate viscosity. This is because, with the increase in viscosity, the substrate relaxes slowly and thus, the effective substrate stiffness increases. Once the substrate elastic stiffness or viscosity is increased significantly, the individual bond force becomes very high, which in the absence of any adhesion reinforcement mechanism only serves to weaken the bonds, which start to fail resulting in lower traction force and higher retrograde velocity. On the contrary, in the case of large binding affinity, *α*, from Figure 4c and 4d, we observe that, at a lower substrate viscosity, increasing substrate elastic stiffness yields a monotonic increase of cell traction force and monotonic decrease of retrograde flow. Same, monotonic behaviour of traction force and retrograde flow is observed when substrate viscosity is increased, keeping the elastic stiffness low. In the presence of high substrate viscosity or elastic stiffness, the force loading rate increases rapidly that in turn increases the effective binding rate and reinforces the adhesion further, and thus the traction force goes up. However, beyond a threshold value of substrate viscosity, the cell loses its sensitivity towards variation in substrate elasticity resulting in perpetually high traction force and low retrograde velocity, and vice-versa.

**Figure 4.**
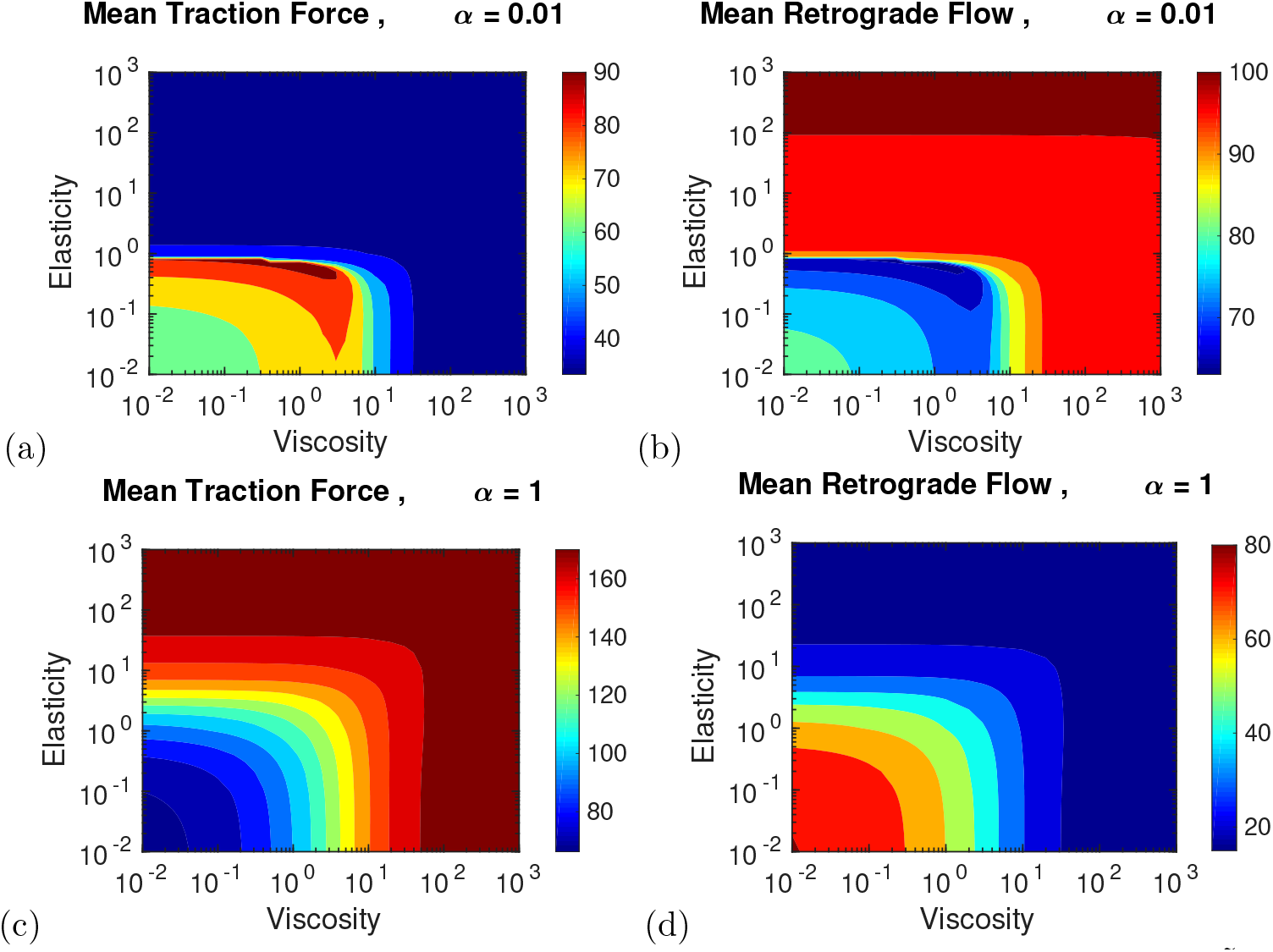
Contour plots representing the mean value of (a) the traction force, 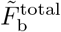, and (b) the retrograde velocity, 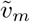, with varying substrate elasticity, 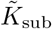, and substrate viscosity, 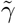, when loading rate dependent adhesion reinforcement is low, *α* = 0.01 (exhibiting biphasic behaviour). Contour plots of mean value of (c) the traction force, (d) the retrograde velocity when loading rate dependent reinforcement is high, *α* = 1 (exhibiting monotonic behaviour).

## Discussion

We have developed a theoretical model consisting of coupled reaction-diffusion equations, which involve force loading rate dependent binding of cell-substrate adhesions at the leading edge of a cell. Owing to the cellular ability to sense and regulate the loading rate sensitive ligand-receptor binding affinity, our model could unravel the puzzling observations of both the biphasic and monotonic relationship of actin retrograde flow and cell traction force with increasing substrate rigidity. Cells that do not have a mechanism to modulate the level of formation of ligand-receptor bonds with the growing force loading rate display a biphasic behaviour, while cells that are able to adapt to increasing loading rate through an adhesion reinforcement mechanism by increasing binding rate display a monotonic increase of traction force and a monotonic decrease of retrograde flow with increasing substrate rigidity. Our study further elucidates the effect of mechanical properties of the substrate, its rigidity and the stress relaxation mechanisms on these nonlinear cellular responses. Besides, it also predicts the loss of cell sensitivity towards the variation of substrate elastic stiffness in case of high substrate viscosity and vice-versa. This has been observed in both the cases of biphasic and monotonic behaviours; although, the nature of cellular response turns out to be quite different. In the case of exhibiting biphasic behaviour, cellular ligand-receptor binding affinity is insensitive to the force loading rate, the bonds quickly dissociate under the high loading rate without getting the opportunity to form more adhesion bonds; this leads to low traction force and high retrograde flow velocity on stiff substrates irrespective of variation in substrate viscosity and vice versa. On the other hand, when the binding affinity increases with increasing loading rate, it allows the formation of more bonds on stiffer substrates that are able to form stable adhesions to support the transmission of cytoskeletal forces, thus leading to the high traction force and low retrograde flow velocity which remains invariant irrespective of changes in substrate viscosity and vice versa. Moreover, we have also investigated the dynamics of cell-substrate adhesions demonstrating both catch bond and slip bond behaviours (presented in the supplementary), and we find a similar biphasic and monotonic relationship with substrate rigidity for both cases. This suggests that the existence of either biphasic or the monotonic relationship is not dependent on the nature of catch or slip bonds. As our study reveals, these nonlinear behaviours can be attributed to the quantitative measure of the cell’s ability to sense and respond to the variation in force loading rate; and that further explains different responses of biphasic versus monotonic behaviour across diverse cell types. Thus, our work could help in throwing light on how cells perceive and react to substrate mechanical properties and envisage to pave the way to further extended studies in areas where cell-substrate mechanical interactions play a central role in regulating cellular behaviours and functions.

## Acknowledgments

The authors acknowledge the financial support from Science and Engineering Research Board (SERB), Grant No. SR/FTP/PS-105/2013, Department of Science and Technology (DST), India.

## Competing interests

The authors declare no competing interests.

## Supplementary Information

### Introduction

In this paper, we have presented a model for cell-substrate adhesions at the leading edge of a cell, displaying both the biphasic and monotonic behaviour of retrograde flow and traction force with increasing substrate rigidity. These nonlinear behaviours could be explained based on the ability of cells to sense the variation of the force loading rate and consequently modulate the ligand-receptors binding affinity to reinforce adhesions in response to fast-growing forces. In the paper, we presented the results considering the ligand-receptor adhesion bonds as catch bonds. However, the results of this model hold good for both catch bonds as well as slip bonds behaviours.

As discussed in the paper, corresponding dissociation rates for catch bonds and slip bonds are expressed as,

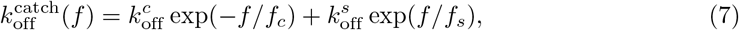

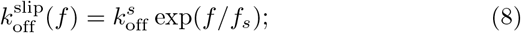

with 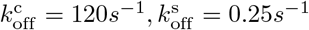, *f*_s_ = 1*pN* and *f*_c_ = 0.5*pN* [37]. Figure 5 shows the dissociation rate for catch bonds and slip bonds with increasing bond force (in the scaled unit presented in the paper).

**Figure 5.**
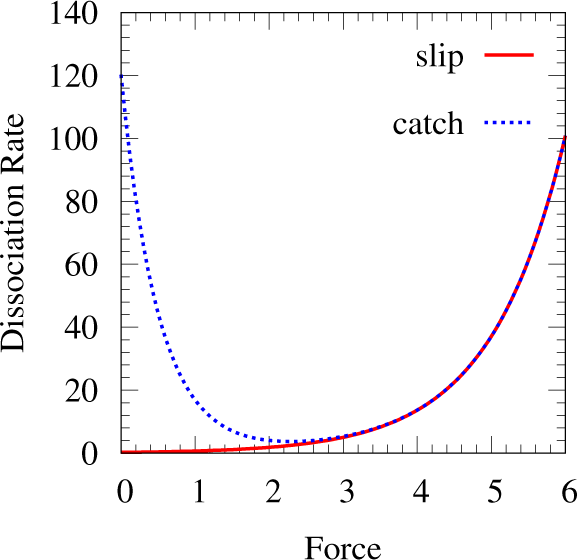
Plots of dissociation rate for catch bonds (blue doted curve) and slip bonds (solid red curve) as a function of bond force.

### Stick-slip motion of actin retrograde flow considering catch bonds behaviour

Figures 6(a)-(d) show the time evolution of retrograde flow velocity, 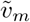, on soft and stiff substrates for two different values of force loading rate dependent binding affinity, *α*; (other parameters are kept at the same value as in the paper). Corresponding traction force, 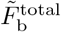, plots are shown in the paper in Figure 2. As expected on a soft substrate, there is not much noticeable change in the time evolution of retrograde flow with the variation in *α* as the force loading rate on a soft substrate is quite low. However, as the substrate stiffness is increased, one does observe a distinct difference in the time evolution as the higher loading rate triggers the binding of ligand-receptor bonds at a higher rate and strengthens the adhesions leading to much more slowing down of retrograde velocity and also increase in the stick-slip cycle duration.

**Figure 6.**
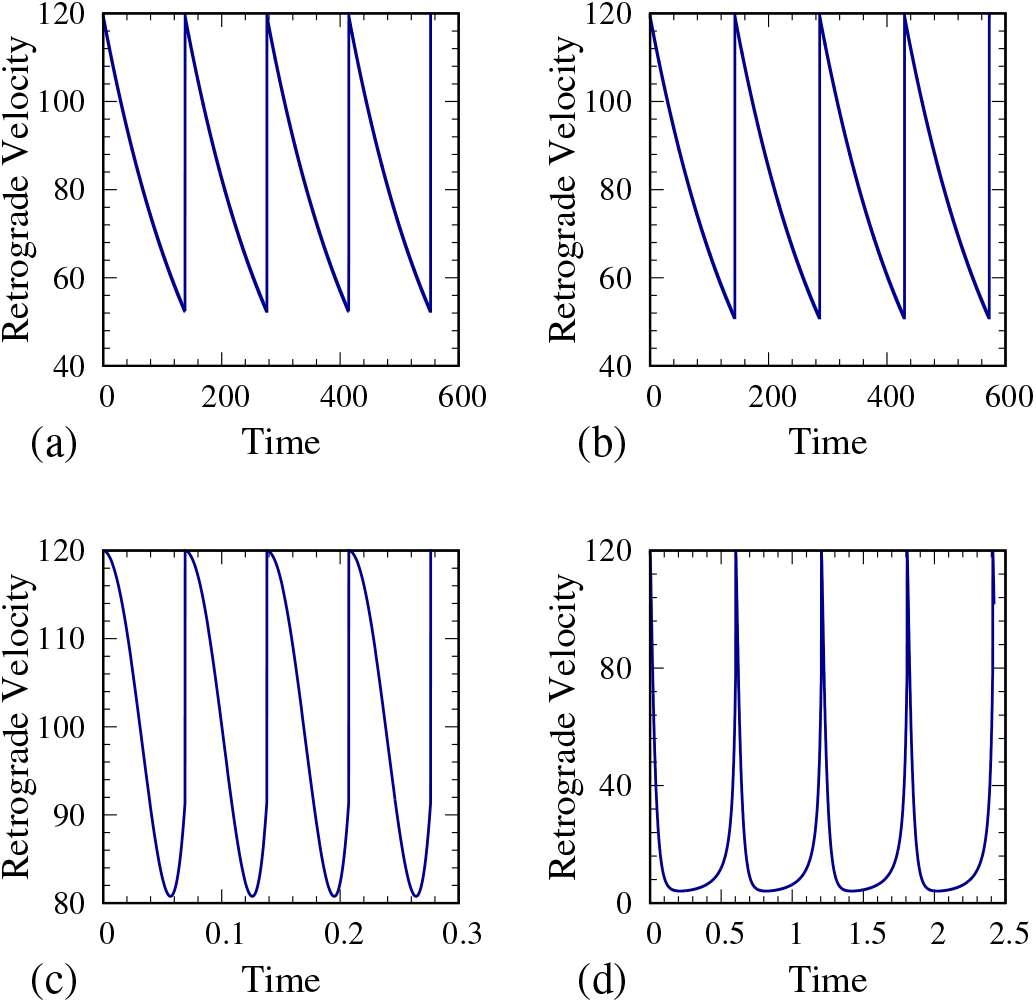
Stick-slip motion for catch bonds. Time evolution of retrograde flow velocity, 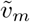, on soft and stiff substrates for two different values of the binding affinity, *α*, with (a) 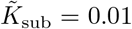, *α* = 0.01; (b) 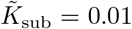, *α* =1; (c) 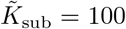, *α* = 0.01; and (d) 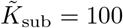, *α* = 1.

### Stick-slip motion for slip bonds

Figures 7 and 8 show the time evolution of the traction force, 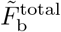 and actin retrograde flow velocity, 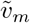, for slip bonds on soft and stiff substrate with low and high values of force loading rate dependent binding affinity, *α*. The stick-slip motion, as we can see, is very similar to their catch bond counterparts.

**Figure 7.**
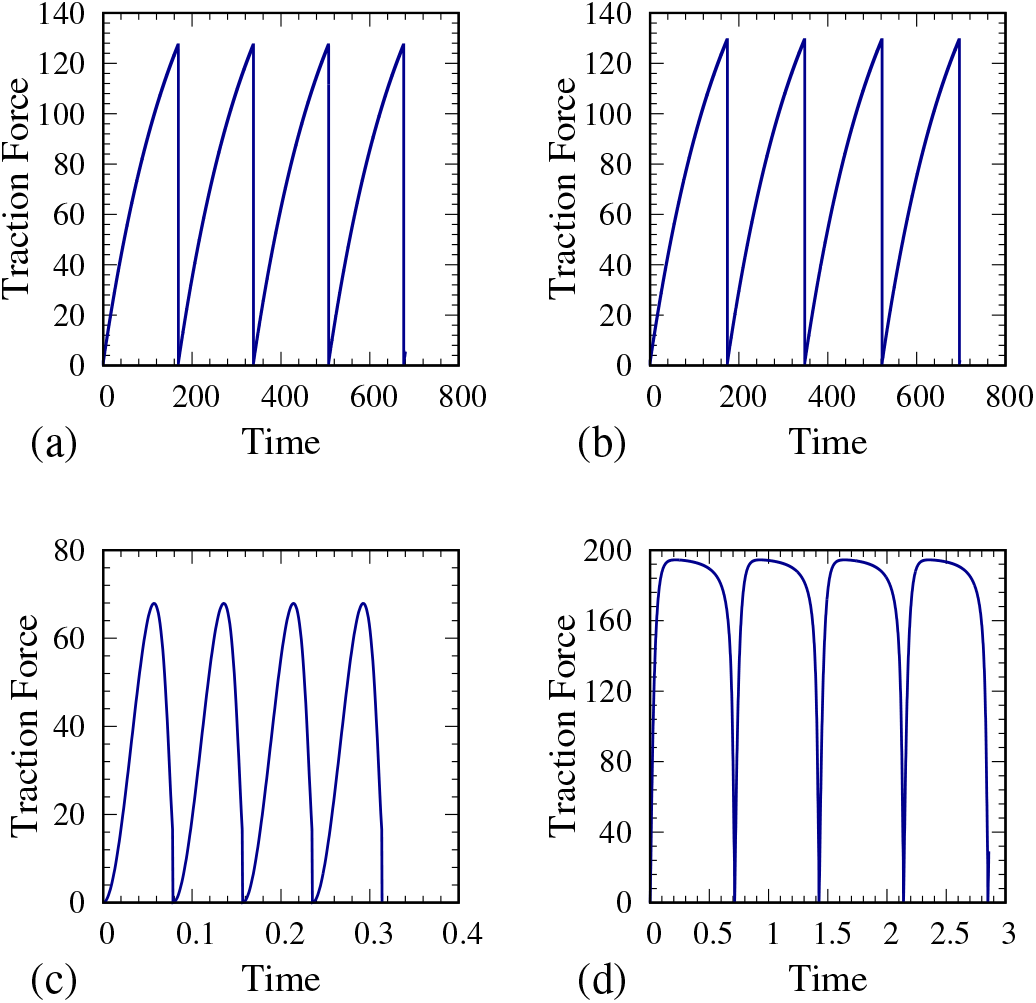
Stick-slip motion for slip bonds. Time evolution of traction force, 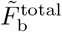, on soft and stiff substrates, with two values of the binding factor, *α*, keeping (a)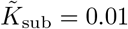, *α* = 0.01; (b) 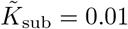, *α* =1; (c) 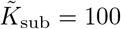, *α* = 0.01; and (d) 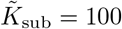, *α* = 1.

**Figure 8.**
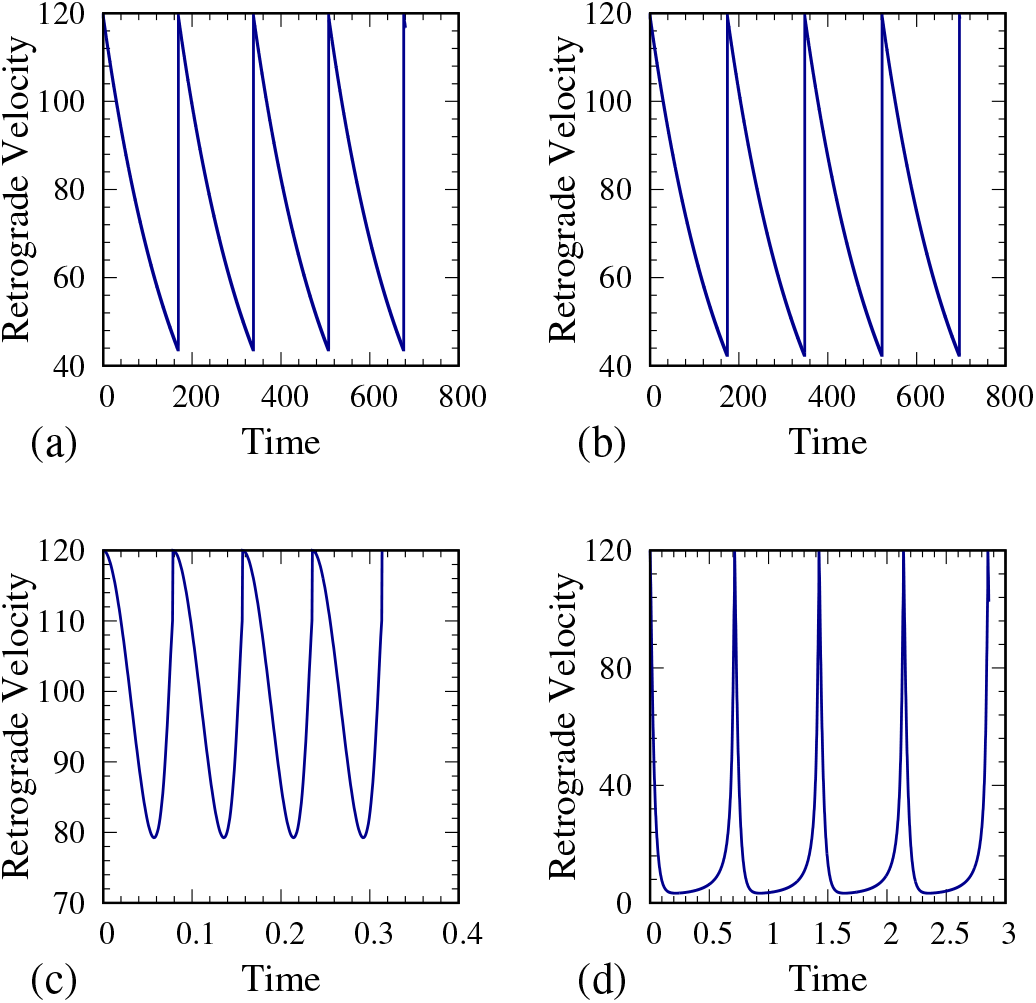
Stick-slip motion for slip bonds. Time evolution of retrograde flow velocity, 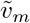, on soft and stiff substrates, with two values of the binding factor, a, with (a) 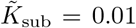, *α* = 0.01; (b) 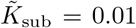, *α* =1; (c) 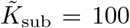, *α* = 0.01; and (d) 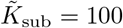, *α* = 1.

### Effect of viscoelastic properties of the substrate

Considering slip bonds behaviour, Figures 9(a)-(b) show the average value of cell traction force, 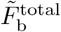, and the average retrograde velocity, 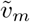, with increasing substrate elastic stiffness, *K*_sub_, keeping the substrate viscosity, 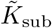, constant, for different values of force loading rate dependent binding affinity, *α*. As observed in the case of catch bonds, here for slip bonds, with the increase in the binding factor, *α* = 0.01, 0. 1, 1, we also see a clear transition from biphasic to the monotonic relationship between the traction force and the retrograde flow with the substrate stiffness.

**Figure 9.**
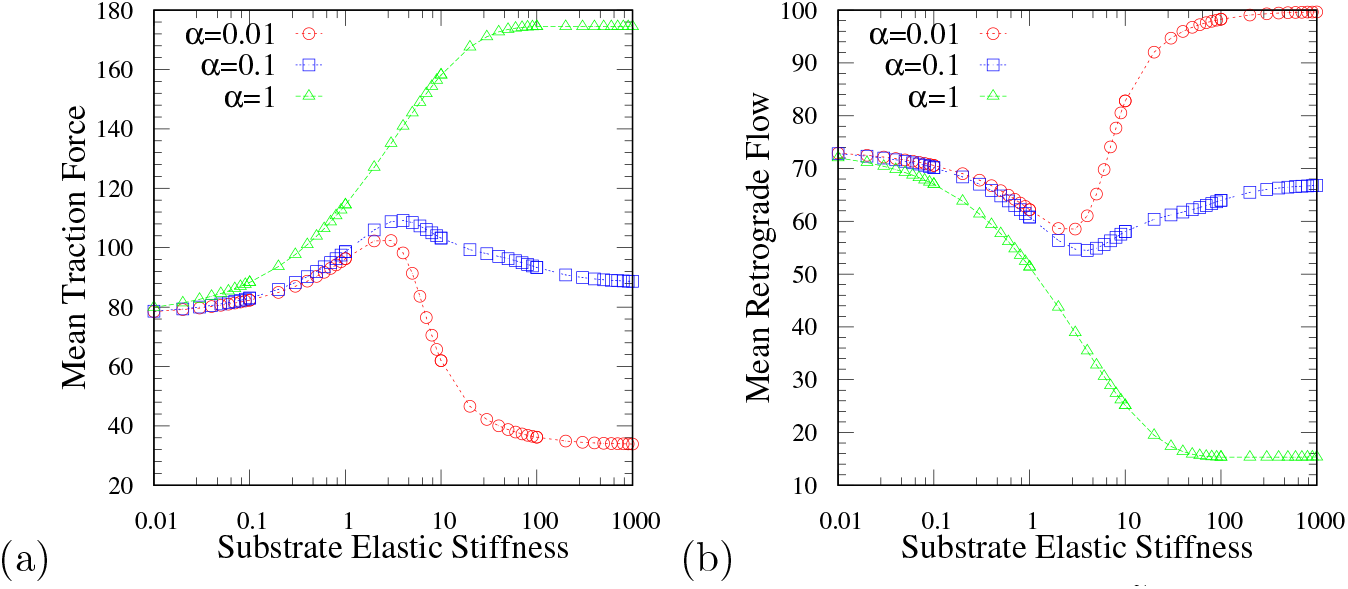
For slip bonds, average value of (a) the traction force, 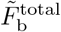, and (b) the retrograde flow velocity, 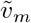, exhibiting the transition from the biphasic to the monotonic relationship of traction force and retrograde velocity with increasing substrate elastic stiffness, 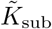, with varying values of the binding factor, *α* = 0.01, 0.1, and 1; (substrate viscosity is kept constant at 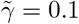).

We have further, investigated whether the duration of stick-slip cycles gets affected by the slip bonds and the catch bonds behaviours. Thus, we have numerically computed the cycle duration for both the cases with varying substrate stiffness. Moreover, we have obtained an analytical expression of the stick-slip cycle duration, *τ*_c_, from our model through a rudimentary calculations given as,

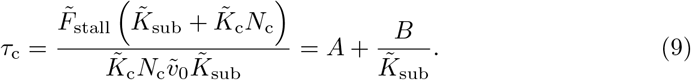

Where the *A* and *B* are given by 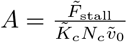 and 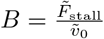, where 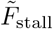 is the scaled stall force, 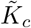 is the clutch stiffness, *N*_c_ is the number of bound receptors and *υ*_0_ is the unloaded retrograde velocity [30]. From Figure 10, it is quite evident that the dependence of the stick-slip cycle can be predicted from the Eq. 9 and the duration is not much altered by the slip or the catch bond behaviours. Further, the qualitative dependence of the stick-slip cycle duration on substrate elastic modulus also remains unaltered in both the cases of low and high loading rate dependent reinforcement factor which show the biphasic and the monotonic behaviour of actin retrograde flow with substrate elastic stiffness.

**Figure 10.**
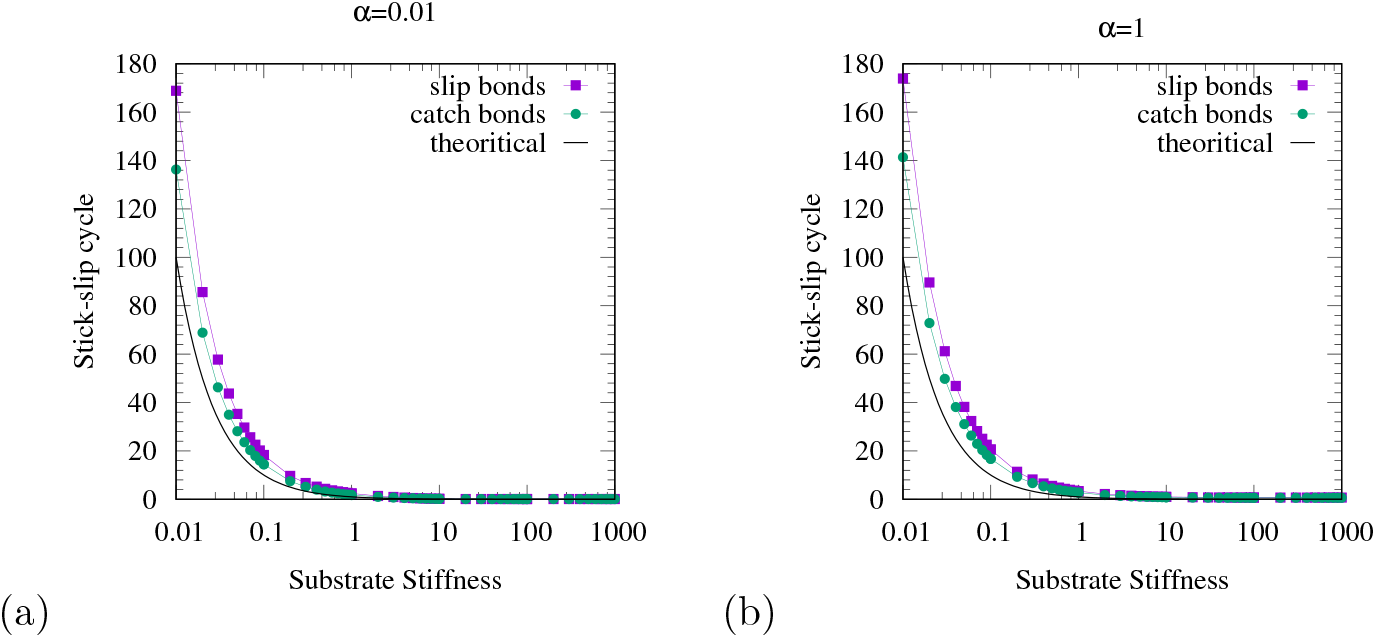
Variation of stick-slip cycle duration on substrate stiffness for catch bonds and slip bonds along with analytical prediction, Eq. 9, for both the cases exhibiting biphasic and monotonic behaviour for (a) low and (b) high values of loading rate dependent reinforcement factor, *α*, respectively.

Further, from the contour plots of Figure 11, we see how the substrate mechanical properties, both elasticity, and viscosity, affect the biphasic and monotonic responses in the case of slip bonds. Figures 11(a) and 11(b) show when the binding factor is low, *α* = 0.01, there exists a biphasic relationship of traction force and retrograde velocity with substrate elasticity and viscosity, with the presence of an optimal substrate viscosity and elasticity corresponding to maximal traction force and minimal retrograde flow. On the other hand, for a higher value of *α* = 1.0, *i.e.*, in the presence of a strong loading rate dependent binding affinity, we find a monotonic relationship between traction force, retrograde flow, and substrate rigidity. These behaviours remain the same as observed for catch bonds that have been presented in the manuscript. All of these goes to show that the emergence of biphasic and monotonic behaviours of the cell traction force and the retrograde flow with the substrate rigidity is not affected by the dissociation rates being slip type or catch-slip type; instead, it is determined by the ability of cells to sense force loading rates and adapt to the growing force as demonstrated by our study.

**Figure 11.**
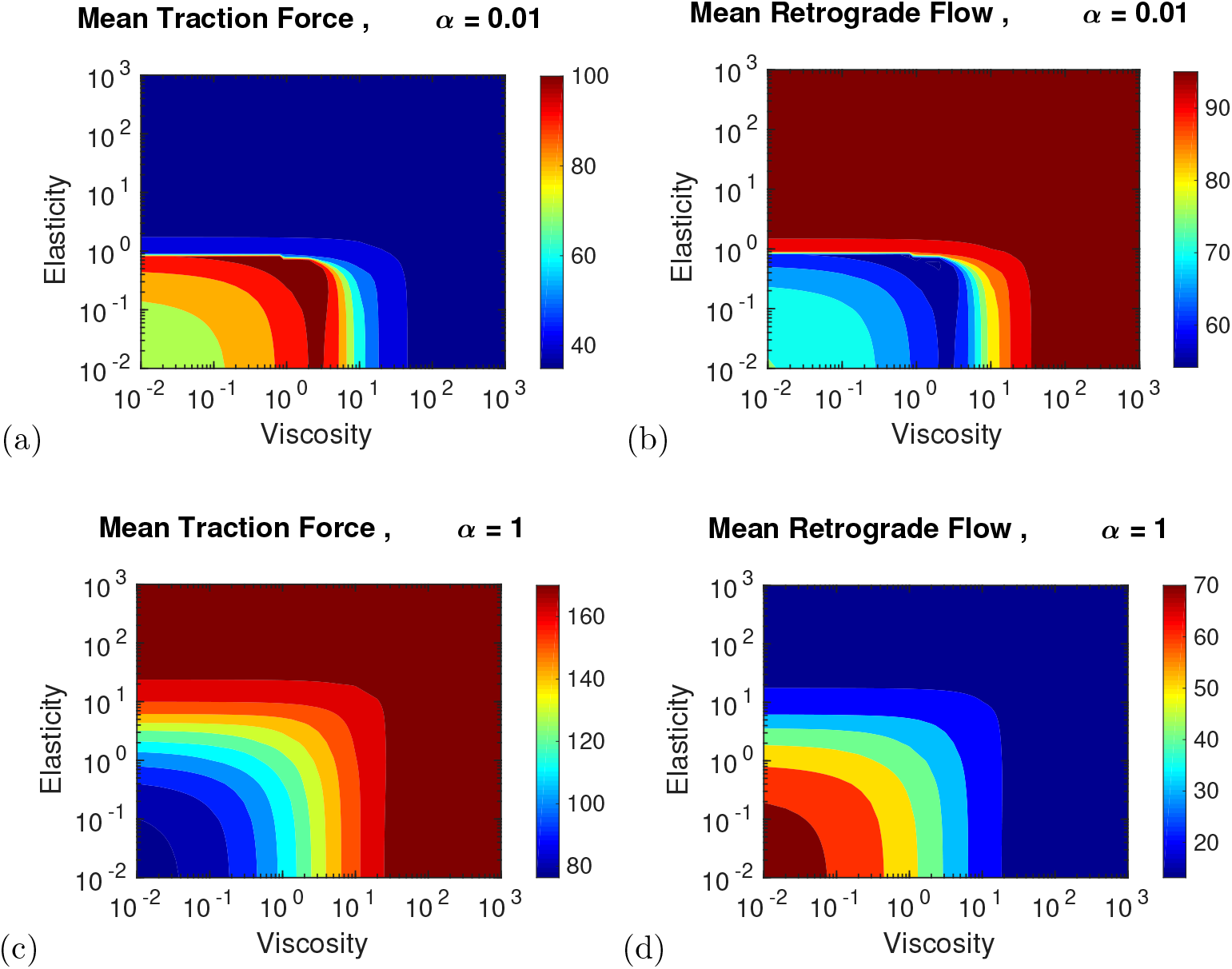
In the case of slip bonds, contour plots representing the mean value of (a) the traction force, 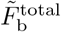, and (b) the retrograde velocity, 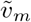, with substrate elasticity and substrate viscosity, when loading rate dependent binding factor is low, *α* = 0.01, exhibiting biphasic relationship. Contour plots of the mean value of (c) the traction force and (d) the retrograde velocity when loading rate dependent binding affinity is high, *α* = 1, showing emergence of monotonic relationship.

## Notes

### Competing Interest Statement

The authors have declared no competing interest.

